# Removing an S-layer enables endolysin-derived probes to detect *Paenibacillus thiaminolyticus* in mixed bacterial populations

**DOI:** 10.64898/2026.09.23.753664

**Authors:** Sarah K. Szwed, Danielle S. McGrath, Morgan C. Serbagi, Chad W. Euler, Joseph N. Paulson, Sarah U. Morton, Jessica E. Ericson, Marwan Osman, James R. Broach, Steven J. Schiff, Vincent A. Fischetti, Edmondo Campisi

## Abstract

The emerging neonatal bacterial pathogen *Paenibacillus thiaminolyticus* causes sepsis and postinfectious hydrocephalus, yet few tools are available to study or specifically detect it. We developed fluorescent probes based on cell wall-binding domains (CBDs) from bacteriophage-derived cell-wall hydrolases. CBDs can target structurally conserved cell-wall glycans, making them attractive affinity reagents for bacterial detection. None of the 11 probes we designed labeled intact cells under standard conditions, and we identified a surface layer (S-layer) that prevented access to their cell-wall targets. Brief acidic treatment (pH 2.0) released abundant high-molecular-mass proteins from the cell surface and enabled labeling by five distinct CBD probes, each containing a single C-terminal S-layer homology domain. Mass spectrometry identified the main released protein as an ortholog of the *P. alvei* SpaA S-layer protein, which we designate SlsA. Across 26 surveyed *P. thiaminolyticus* genomes, we identified four distinct SlsA variants with 58–79% pairwise amino acid identity, encoded within a conserved cluster homologous to the *P. alvei* S-layer-associated locus. After either the acidic treatment or a quick flame fixation, the lead probe mGL-BDPt9 labeled six *P. thiaminolyticus* strains, including three Ugandan clinical isolates, as well as a *P. dendritiformis* clinical isolate from a US infant, and three additional *Paenibacillus* species. mGL-BDPt9 also distinguished *P. thiaminolyticus* from five gram-positive sepsis pathogens in pairwise mixtures and detected it in a five-species suspension. Together, these findings establish the presence of an S-layer in *P. thiaminolyticus* and provide a rapid fluorescent labeling strategy for multiple *Paenibacillus* pathogens associated with neonatal paenibacilliosis.

**IMPORTANCE:** *Paenibacillus* species are increasingly recognized as causes of devastating neonatal infections, but little is known about the surface structures of these pathogens or how those structures affect direct detection. Here, we identify an S-layer in *P. thiaminolyticus*, revealing a major cell-surface component that may contribute to pathogenesis, since S-layers mediate host interactions and contribute to immune evasion and virulence in other bacterial pathogens. We also show that the S-layer can hide conserved cell-wall targets and can be rapidly removed with a simple one-minute treatment that is readily incorporated into labeling protocols. Probes built from the binding domains of prophage-encoded enzymes labeled clinical isolates of two *Paenibacillus* species associated with infant disease and distinguished *P. thiaminolyticus* within mixed bacterial suspensions. S-layers are common among bacterial pathogens, and similar removal steps may expose additional cell-wall targets for affinity-probe development in those organisms.

## INTRODUCTION

Every year, 6.9 million neonates in sub-Saharan Africa, South Asia, and Latin America are treated for possible severe bacterial infection, with 680,000 associated deaths (1). Treatment is generally empirical and broad-spectrum, since the diagnosis is primarily clinical, and identification of a causative organism depends on slow, low-sensitivity culture (2). Sepsis and meningitis are the main presentations, and survivors of the latter have an increased risk of impaired neurodevelopment and postinfectious hydrocephalus (PIH) (3).

Neonatal infection causes 60% of infant hydrocephalus in Uganda (4), where a third of infants treated for PIH die within five years, and a third of those who survive suffer from severe neurodevelopmental impairment (5). The bacterial genus most frequently identified in cerebrospinal fluid from infants with PIH is *Paenibacillus* (3). Among 631 neonates with clinical sepsis, 6% tested positive for *Paenibacillus*, of whom 70% were *P. thiaminolyticus*. Thirty percent of these neonates died or developed hydrocephalus or neurodevelopmental impairment within one year, compared to 13% of neonates with sepsis from other causes (6). Two of three isolates from infants with PIH were resistant to ampicillin, included in the WHO empirical regimen, and none was vancomycin-susceptible (6). Following the Ugandan reports, the same presentation with cystic encephalomalacia has been described in infants across multiple U.S. states (7, 8). Although *P. thiaminolyticus* is the predominant species among the Ugandan cases, other members of the genus have also been reported to cause human disease, including *P. dendritiformis* and *P. alvei* (7, 9, 10).

Infants reach the hospital once hydrocephalus has developed following meningitis, when treatment is palliative, making earlier detection critical to an effective intervention. Further complicating detection, *Paenibacillus* stains gram-variable, and recovery of the Ugandan isolates required anaerobic culturing conditions and blood-supplemented media, not routinely used in neonatal sepsis diagnostic protocols. In both reported U.S. infant cases, MALDI-TOF mass spectrometry called *P. thiaminolyticus* and sequencing returned *P. dendritiformis* (8). No direct molecular reagent for detecting *Paenibacillus* is currently available.

Bacterial and phage genomes encode a diverse functional class of cell-wall hydrolases. Among them, bacteriophage endolysins, or lysins, degrade the host cell wall to release progeny virions and are being developed as a novel class of antibacterials because they rapidly kill gram-positive bacteria when applied exogenously (11, 12). Most lysins are modular, comprising an enzymatically active domain (EAD) and a cell wall-binding domain (CBD) that recognizes bacterial surface ligands with high affinity. CBDs can be functionally expressed in isolation and, when fluorescently labeled, serve as affinity reagents that are derived directly from genomic sequences and target conserved surface carbohydrates, with specificity ranging from a single strain to an entire genus. CBD-based probes have been developed for several bacterial pathogens (13, 14), including the plant-associated *P. polymyxa* and the honeybee pathogen *P. larvae* (15, 16), but none for *P. thiaminolyticus* or the other species associated with neonatal PIH.

In addition to the canonical cell wall components, some bacteria have an external proteinaceous paracrystalline monolayer, the S-layer, or surface layer, that contributes to structural maintenance while remaining dynamic during cell division (17). In many gram-positive organisms, the S-layer is attached to the cell wall through non-covalent interactions between an S-layer homology (SLH) domain and a peptidoglycan-linked secondary cell wall polymer (SCWP), typically one including pyruvylated sugar residues (18, 19). Some SLH domains also recognize peptidoglycan directly, as described for *P. alvei* (20). To develop a diagnostic probe for *Paenibacillus*, we identified eleven cell-wall hydrolase candidates, including phage endolysins, from publicly available genome assemblies classified as *P. thiaminolyticus*, and produced fluorescent fusions of their binding domains alone. After brief pre-treatment to disrupt the envelope, five probes, each carrying a single C-terminal SLH domain, labeled the organism clearly with well-defined outlines. Finally, we identified the barrier as an S-layer that occluded their cell-wall targets (21, 22), and showed that its removal enables probe-mediated labeling of reference strains and clinical isolates and detection of the pathogen in mixed bacterial populations.

## MATERIALS AND METHODS

### Bacterial strains and growth conditions

All the strains used in this study are listed in Table S4. *Paenibacillus* strains were grown in Todd-Hewitt broth, other gram-positive species in Brain Heart Infusion, and *Escherichia coli* in Luria-Bertani, in broth (with shaking) or on solid medium containing 15 g/L agar. Todd-Hewitt broth was supplemented with 0.1% (wt/vol) Tween 80 for *P. macerans* ATCC 8509 and *P. larvae* ATCC 9545. The clinical isolates used in this study were obtained from previously described collections and were examined as de-identified archived bacterial cultures, with no access to identifiable patient information.

### In-silico identification of candidate cell-wall hydrolases

Twenty-six *P. thiaminolyticus* genome assemblies publicly available at the time of analysis (Table S2) were downloaded from NCBI Datasets (23), and their predicted proteins were annotated with InterProScan (24, 25), reducing overlapping domain calls to one per region. Candidate cell-wall hydrolases were selected on their predicted catalytic and cell-wall-binding domains. For each candidate, we manually inspected annotated neighboring genes in the deposited assemblies and scored it as a phage endolysin if we identified a putative holin within 5 ORFs from the candidate. For lysin candidates encoded on complete genomes, we tested prophage origin with PHASTEST (26). Redundant candidates were removed by collapsing groups sharing ≥75% identity across the full-length protein (pairwise MAFFT alignment (27)) to a single representative, and eleven unique sequences, LysPt1 to LysPt11, were carried forward for binding-domain cloning (Table S1).

### Fluorescent probe design and expression

The putative binding domain of each candidate (Table S1) was amplified from plasmids containing the full parental coding sequence and cloned by in vivo assembly into a custom-modified pET14b vector in frame downstream of an N-terminal avi-tag–6×His–monomeric GreenLantern–(GGGGS)₃ fusion, giving mGL-BDPt1 to mGL-BDPt11. A *Clostridium perfringens*-specific binding domain was cloned into the same vector, with the same architecture, to generate mGL-BDCp17. Its characterization and binding specificity are demonstrated in [thesis, doi 10.48496/9pw7-h927]. Constructs were expressed in *E. coli* BL21-AI and used either as clarified sonicated lysate or after Ni-NTA purification. Construction, expression and purification are described in Supplementary Methods.

### Envelope-disrupting cell pretreatment

Strains were streaked on Todd-Hewitt agar plates and grown overnight at 37 °C. Biomass was resuspended from the plate into Buffer A, vortexed to homogeneity, and adjusted to OD₆₀₀ 0.6. Two 1 mL aliquots were pelleted (10,000 × g, 2 min) and washed once in 1 mL of Buffer A, and each pellet was resuspended in 100 µL of either Buffer A or 0.2 M glycine-HCl pH 2.0 and incubated for 1 min at room temperature. Suspensions were neutralized with 1 mL of Buffer A, re-pelleted, and resuspended in 100 µL of Buffer A. The best treatment condition was selected from a pH and time series scored at the microscope (Table S3).

### Labeling and mounting

Cells were labeled by one of two routes. In solution, 10 µL of cell suspension was mixed with 10 µL of probe, incubated 10 min at room temperature, diluted with 500 µL of Buffer A (50 mM HEPES, 140 mM NaCl, pH 7.4), pelleted (10,000 × g, 5 min), resuspended in 20 µL of Buffer A, and 2 µL was spotted and covered with a No. 1.5 coverslip. On the slide, cells were diluted 1:1 in Buffer A, 2 µL was spotted and flame-fixed, 3 µL of probe was applied, the excess was drawn off, and 3 µL of Buffer A was added before covering. For 100× and 60× imaging, cells were counterstained with BacLight Red, fixed in 0.5% paraformaldehyde (1 h, 37 °C), washed, labeled with purified mGL-BDPt9 at 0.3 µg/µL for 10 min at room temperature, washed, and mounted on a 1% agarose pad in water sealed with VALAP. Pairwise co-mixtures were normalized by optical density, combined, and carried through this route.

### Fluorescence microscopy

Imaging at 40× used a Nikon Eclipse E400 with a Plan 40×/0.65 Ph2 DL objective and a DMK 23UX174 camera (0.1465 µm/pixel, 16-bit), under phase contrast (50 ms, gain 1.5, gamma 1.30) and epifluorescence (500 ms, gain 1.9, gamma 0.75) in the FITC channel. Settings were identical for every field reported. Fixed cells were imaged on a DeltaVision system with an sCMOS camera, the reference strain and pairwise co-mixtures with an Olympus 100× NA 1.40 objective (0.0646 µm/pixel) and the clinical isolates at 60× (0.1074 µm/pixel), over 12 to 14 optical sections (14 to 22 for the clinical isolates), with emission collected at 525 and 632 nm. Stacks were bleach-corrected, deconvolved against a measured optical transfer function and projected by maximum intensity. The operator examined each field and scored it positive or negative against background.

### Whole-cell binding assay

*P. thiaminolyticus* ATCC 13023 at OD₆₀₀ 1 was treated with or without 0.2 M glycine-HCl pH 2 for 5 min, washed, incubated with clarified mGL-BDPt7 lysate for 20 min at room temperature, and then washed twice more. Total, supernatant and pellet fractions were taken after those washes and resolved by SDS-PAGE. Adapted from (22).

### SDS-PAGE

Samples were prepared in NuPAGE LDS sample buffer with reducing agent, heated 5 min at 95 °C, and resolved on NuPAGE 4–12% Bis-Tris gels at 200 V in MOPS running buffer for the fractionation shown in Fig. 3A and MES for that in Fig. 3B, against the Novex Sharp pre-stained standard, then stained with SimplyBlue SafeStain.

### Protein identification by mass spectrometry

Bands representing the surface proteins extracted from the surface of *P. thiaminolyticus* ATCC 13023 with 0.2 M glycine-HCl pH 2 for 1 min were excised from lane 6 of the MOPS-buffered gel shown in Fig. 3A, then reduced, alkylated, and digested with trypsin overnight. One third of each digest was analyzed by LC-MS/MS (Orbitrap Ascend, ES902 column, HCD fragmentation). Spectra were searched with Mascot (28) in Proteome Discoverer 1.4 (Thermo Fisher Scientific) against UniProt proteome UP000315377 (*P. thiaminolyticus* NRRL B-4156) supplemented with additional S-layer protein sequences and a common-contaminant background, using trypsin with up to three missed cleavages, 10 ppm precursor and 20 mmu fragment tolerance, fixed carbamidomethylation of cysteine, and variable oxidation of methionine and N-terminal acetylation. Peptides were filtered at high confidence, rank 1, and proteins grouped by strict maximum parsimony. Glycosylation was assessed in glycan mode in PEAKS Studio v12.5 (Bioinformatics Solutions Inc.) (29) and by inspection of HCD spectra for glycan oxonium ions.

### SLH-domain annotation

S-layer homology domains were identified in the annotated *P. thiaminolyticus* NRRL B-4156 genome (GenBank NZ_CP041405.1, 5,702 coding sequences) with HMMER v3.4 (30) and the Pfam model PF00395.26 (31), calling hits at the curated gathering thresholds and reporting domain boundaries at a per-domain threshold of E ≤ 1.

### SCWP and S-layer locus biogenesis comparison

S-layer biosynthesis loci were retrieved from RefSeq genome records with flanking sequence extending to *fabZ* and *metK*, and aligned in Geneious Prime 2026.1.2 using MAFFT v7.490 (27) under the L-INS-i strategy with the PAM200 / k = 2 scoring matrix and default gap parameters.

### Image and figure preparation

Gel photographs were processed for figures in Fiji (ImageJ 1.54p) (32) by extracting the red channel, applying an adjustment to the levels, and cropping. Micrograph windowing, registration and cropping were performed in the same software. Only linear adjustments applied uniformly to the whole image were used. Figures were composed in Adobe Illustrator 27.0.1.

### Data availability

Assembly accessions for the genomes surveyed are given in Table S2 and for the candidate panel in Table S1. Original image files are retained and available on request. Part of this work is reported in a doctoral thesis (https://doi.org/10.48496/9pw7-h927).

## RESULTS

### Five CBDs from a panel of eleven Paenibacillus thiaminolyticus cell-wall hydrolases label the organism after envelope disruption

To identify candidate cell wall-binding domains (CBDs) for *P. thiaminolyticus*, we analyzed the 26 publicly available genome assemblies in the NCBI database at the time of screening. Despite the relatively small number of available genomes, they were a source of multiple prophage regions, with several strains being polylysogenized, as previously reported (33). Maximizing sampling diversity, we selected 11 candidates, LysPt1 to LysPt11, for expression and characterization (Fig. 1).

**Figure 1.**
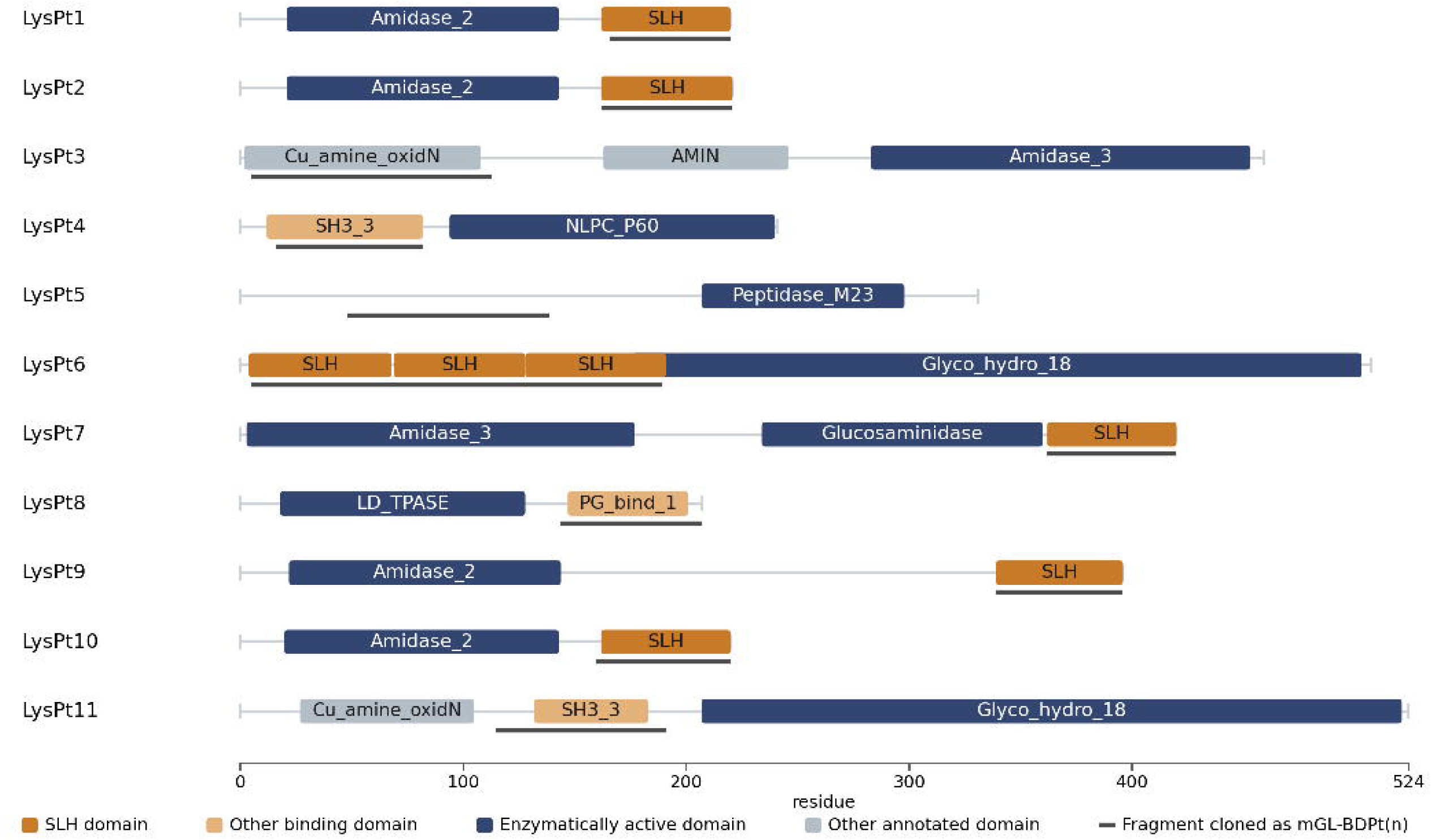
Domain architecture of eleven candidate cell-wall hydrolase of *Paenibacillus thiaminolyticus*. Each candidate is drawn to scale on a base-pair length axis. Domains are colored by class: S-layer homology (SLH) domains; other annotated cell-wall binding domains, comprising SH3_3 and PG_binding_1; enzymatically active domains; and other annotated domains. PG_bind_1 = PG_binding_1. Regions carrying no annotation are drawn as backbone. The line beneath each backbone marks the fragment cloned as that candidate’s mGreenLantern fusion. Labeling results for each fusion are given in Table 1, and the source and properties of each candidate are in Table S1.

**Table 1.** Labeling of eleven mGreenLantern–cell wall binding domain fusions to *Paenibacillus thiaminolyticus* ATCC 13023, before and after envelope disruption.

| Fusion | Binding module cloned | Untreated | Envelope-disrupted |
| --- | --- | --- | --- |
| mGL-BDPt1 | SLH, single C-terminal | — | + |
| mGL-BDPt2 | SLH, single C-terminal | — | + |
| mGL-BDPt3 | Copper amine oxidase, N-terminal domain | — | — |
| mGL-BDPt4 | SH3_3 | — | — |
| mGL-BDPt5 | Unannotated region | — | — |
| mGL-BDPt6 | SLH, N-terminal three-domain module | — | — |
| mGL-BDPt7 | SLH, single C-terminal | — | + |
| mGL-BDPt8 | PG_binding_1 | — | — |
| mGL-BDPt9 | SLH, single C-terminal | — | + |
| mGL-BDPt10 | SLH, single C-terminal | — | + |
| mGL-BDPt11 | SH3_3 | — | — |

The candidates are derived from 6 source assemblies representing strains with clinical and non-clinical origins, including Ugandan cerebrospinal-fluid isolates (Table S1). The panel included representatives of the major cell-wall-hydrolase catalytic classes: N-acetylmuramoyl-L-alanine amidases (Amidase_2 and Amidase_3), an N-acetylglucosaminidase, an NlpC/P60 DL-endopeptidase, an M23 metallopeptidase, an L,D-transpeptidase family (YkuD) enzyme, and glycoside hydrolase family 18 enzymes. Additionally, all selected candidates carried a predicted cell-wall-binding module except LysPt5 (Fig. 1). Specifically, 5 candidates carried an atypical single SLH domain (LysPt1, 2, 7, 9 and 10), one a canonical trimeric SLH module (LysPt6), and the remainder carried SH3, PG_binding_1, AMIN, or copper amine oxidase N-terminal domains. Four candidates, namely LysPt1, LysPt7, LysPt9, and LysPt10, had a putative holin within a local lysis cassette in the near genomic context and were thus classified as phage endolysins. The remaining candidates were non-phage-derived cell-wall hydrolases identified on domain content alone.

S-layer homology (SLH) domains non-covalently anchor surface proteins to peptidoglycan-linked secondary cell wall polymers (SCWPs) and often occur as 3 tandem repeats forming a pseudo-threefold module (34). Guided by the predicted domain boundaries and structural predictions, we expressed one putative binding module from each candidate as a fluorescent probe fused to an mGreenLantern (mGL) reporter in *E. coli*. All 11 probes were successfully expressed in the crude lysate (Fig. S1), with only mGL-BDPt8 showing a visibly depleted soluble fraction. We then screened each probe as clarified crude lysate for its ability to label a suspension of *P. thiaminolyticus* ATCC 13023, but observed no signal by microscopy, except for a few scattered cells, often in clumps and chains (Table 1, Untreated).

S-layers have been reported to prevent cell wall-acting proteins from engaging their targets in other gram-positive bacteria (21, 22), and while none has been characterized for *P. thiaminolyticus*, 3 species in the genus carry one (19, 35, 36). We hypothesized that an analogous layer could explain the result and tested whether disrupting the envelope would allow our probes to label. Therefore, we ran a pilot test in which we subjected a *P. thiaminolyticus* cell suspension to brief on-slide flame fixation and observed that this treatment enabled fluorescent labeling by mGL-BDPt9. To achieve envelope disruption in solution, we then used glycine-HCl, which has successfully been used for extraction in other S-layer-expressing organisms (21, 22). We rescreened all 11 fluorescent CBD probes under these conditions (Table 1, Envelope-disrupted), and 5 of them, namely mGL-BDPt1, mGL-BDPt2, mGL-BDPt7, mGL-BDPt9, and mGL-BDPt10, all carrying a single C-terminal SLH domain (Fig. 1), showed uniform labeling of acid-treated *P. thiaminolyticus* cells (Fig. S2). Notably, except for mGL-BDPt2, all positive probes were phage-derived.

Even after acid treatment, we observed no labeling among the 6 remaining constructs, including mGL-BDPt6 (the N-terminal SLH trimeric module), mGL-BDPt5 (comprising an unannotated region), and the 4 non-SLH domain-containing CBDs (Table 1, Envelope-disrupted). Of the five binders, we chose mGL-BDPt9 for the subsequent characterization experiments.

### A 1-min acid step or a brief flame fixation is sufficient to disrupt the envelope

Having established that envelope disruption permits labeling, we investigated the minimal treatment required for optimal labeling using our purified lead probe mGL-BDPt9, as well as the treatment’s effect on the cells. Labeling improved as pH fell and reached every cell in the field at pH 2 (Fig. 2A to 2D), while extending the treatment from 1 to 5 min did not cause any visible changes in cell-associated fluorescence (Table S3). We therefore chose pH 2 treatment for 1 minute in all subsequent imaging experiments. The flame fixation method was further optimized into a quick-stain protocol that replaces the acid step and its washes with a few seconds of flame exposure (Fig. 2G and 2H).

**Figure 2.**
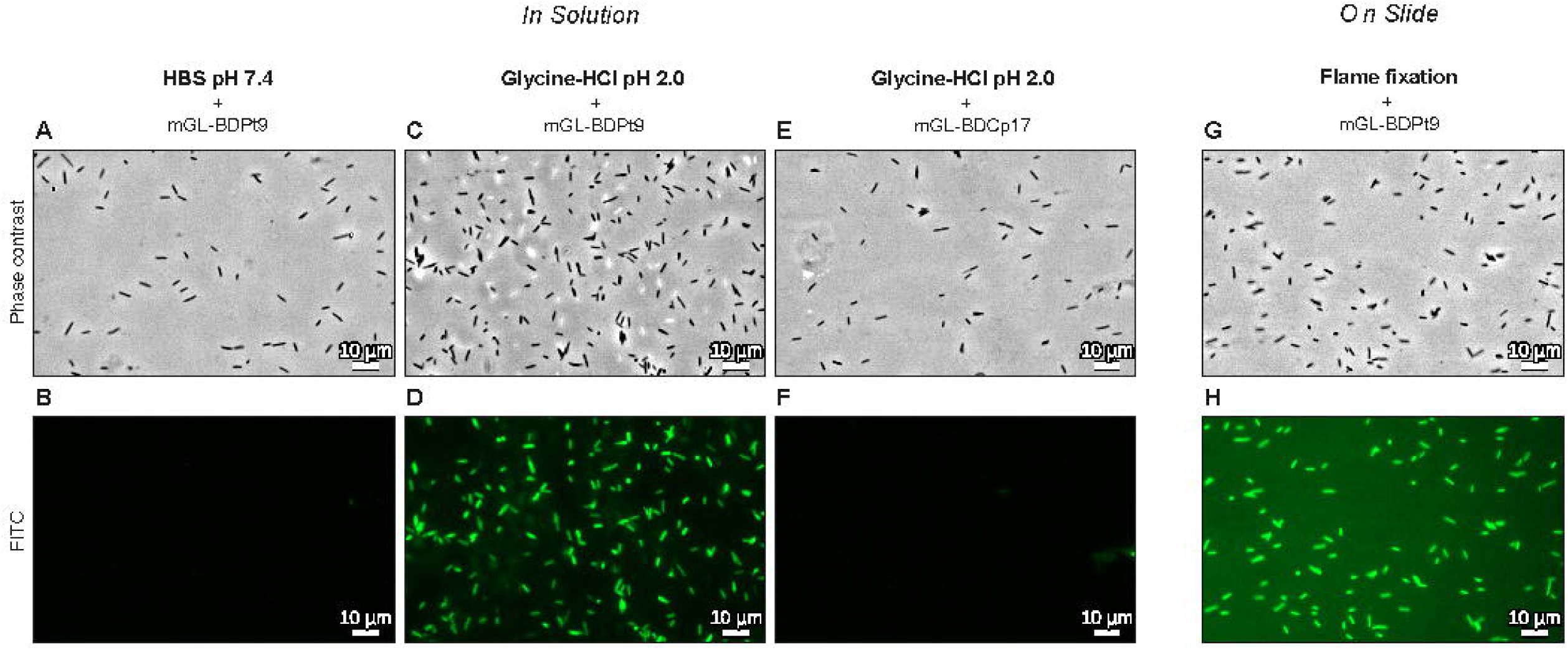
Envelope disruption removes a barrier to probe labeling of *Paenibacillus thiaminolyticus* ATCC 13023. Phase contrast (upper row) and FITC channel (lower row) of the same field. Solution route: (A, B) untreated cells incubated with mGL-BDPt9; (C, D) cells treated with 0.2 M glycine-HCl pH 2.0 for 1 min, then incubated with mGL-BDPt9; (E, F) cells treated identically to C and D but incubated with the heterologous control fusion mGL-BDCp17. On-slide route: (G, H) flame-fixed cells stained on the slide with mGL-BDPt9. Occasional labeling was present in both the untreated and heterologous-control preparations (not pictured here). Panels within a route share one linear display window per channel and were acquired in a single session at identical exposure and gain. The two routes are displayed through separate windows and are not comparable for brightness. Scale bars: 10 µm.

By contrast, acid-treated cells incubated with the heterologous fusion mGL-BDCp17, whose binding module derives from a *Clostridium perfringens* prophage endolysin, showed essentially no labeling, except for the same occasional labeling as untreated preparations (Fig. 2E and 2F).

Interestingly, the pH 2 treatment did not affect gross cell morphology or reduce motility, indicating that, aside from the indirect surface changes we observed, the cells’ integrity was overall preserved. Labeling was robust to variations in culture method, as observed in cells grown in broth at different stages and on agar (Table S3).

### Acid treatment strips an abundant S-layer protein and removes the barrier

The envelope-disrupting treatment we used that enabled labeling is the standard low-pH glycine extraction used to strip S-layers, so we investigated its effect on the cells. We resuspended *P. thiaminolyticus* ATCC 13023 in 0.2 M glycine-HCl pH 2 for 1 min or in neutral control buffer, and resolved the whole suspension, the supernatant and the washed pellet by SDS-PAGE (Fig. 3A).

**Figure 3.**
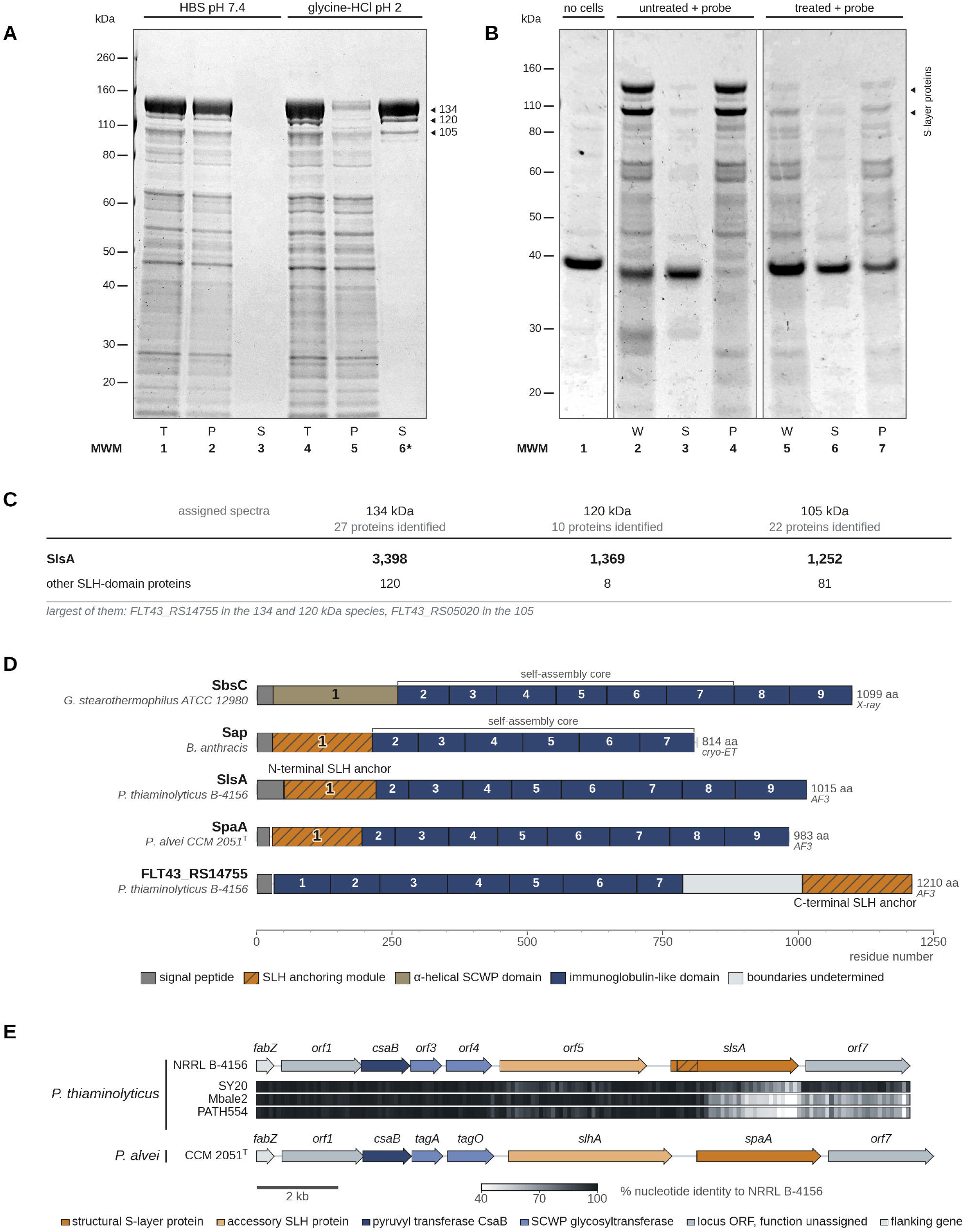
An acid-extractable surface protein of *Paenibacillus thiaminolyticus* is the product of *slsA* and is homologous to characterized Bacillaceae S-layer proteins. (A) Coomassie-stained SDS-PAGE (MOPS). *P. thiaminolyticus* ATCC 13023 cells resuspended in glycine, pH 2, or in neutral control buffer for 1 min. T, whole suspension; S, supernatant after centrifugation; P, pellet washed once and resuspended in neutral buffer. Three high-mass species (134, 120 and 105 kDa) released into the acid supernatant were excised from lane 6 (asterisk) for LC-MS/MS. MWM, molecular weight marker. (B) Coomassie-stained SDS-PAGE (MES). *P. thiaminolyticus* ATCC 13023 cells were subjected to the same acid or neutral-buffer treatment as in A, except that the treatment was extended to 5 minutes, incubated with the cell-wall binding domain probe mGL-BDPt7, washed and the cell fraction was harvested. Lane 1, mGL-BDPt7 control. W, cells + probe (whole reaction); S, supernatant after incubation; P, washed pellet after incubation. High-mass bands abundant in the untreated preparation (lanes 2–4) are depleted after acid treatment (lanes 5–7). Arrowheads mark the two species, later identified as S-layer protein. Following acid extraction, mGL-BDPt7 is retained in the washed cell pellet, consistent with exposure of its cell-wall ligand after removal of the high-molecular-mass surface proteins. (C) LC-MS/MS identification of each excised species, as assigned spectra, searched against the *P. thiaminolyticus* NRRL B-4156 reference proteome. SlsA (S-layer structural protein A; WP_087444342.1) was the dominant match with 93.9 %, 97.0 % and 81.0 % of spectra assigned in the 134, 120 and 105 kDa species, which returned 27, 10 and 22 proteins. The second row pools every other SLH-domain protein identified. Assigned spectra are reported. (D) Predicted domain organization of SlsA and comparison to reference S-layer proteins. SbsC (Geobacillus stearothermophilus, O68840, 1099 aa) and Sap (*Bacillus anthracis*, P49051, 814 aa) are experimental structures; SlsA, SpaA (P. alvei CCM 2051ᵀ, ACN92046.1) and FLT43_RS14755 (WP_087444343.1) are AlphaFold 3 models. Domains are numbered from the N-terminus, counting the anchoring module. Grey, signal peptide; hatched, SLH module; olive, α-helical SCWP domain; blue, β-sandwich domain; pale, boundaries unresolved. Brackets, the self-assembly core of SbsC and Sap. (E) The S-layer and SCWP biosynthesis locus. Upper, *P. thiaminolyticus* NRRL B-4156 (NZ_CP041405.1); lower, P. alvei CCM 2051ᵀ (NZ_AMBZ01000002.1), shown for organization only. Strips, nucleotide identity to NRRL B-4156 in 100-bp windows of a four-genome locus alignment. SY20 (NZ_CP106992.1), Mbale2 (NZ_CP094446.1) and PATH554 (NZ_CP114031.1); darkest = most conserved; blank = fewer than 20 aligned positions.

Acid treatment released abundant high-molecular-mass material from the cell surface into the supernatant. Three species were resolved, at approximately 134, 120 and 105 kDa, which we excised, analyzed by LC-MS/MS and searched against the reference *P. thiaminolyticus* proteome.

In all three electrophoretic species, the top hit was the same protein, with locus tag FLT43_RS14750, which we designate SlsA. SlsA accounted for 93.9%, 97.0% and 81.0% of assigned spectra among the total 27, 10, and 22 proteins returned for the three bands (Fig. 3C). Coverage of the mature sequence nonetheless reached 87.7% (Fig. S3), and was evenly distributed throughout the mature sequence in all three forms, indicating absence of defined truncation events. No glycosylation was detected: a glycan-mode search returned no glycopeptides, and all the commonly *O*-glycosylated acceptor residues in S-layer proteins (Tyr, Thr, and Ser), were detected with unmodified mass. No genome sequence is available for ATCC 13023, but the SlsA peptides recovered from it perfectly match the sequence variant encoded by the type strain NRRL B-4156. The wash also released three further SLH-domain proteins, each a minor component against SlsA, together with one protein that carries no anchoring module (Table S5). Two of these proteins, FLT43_RS14755 and orf7, are encoded within the same locus as slsA (Fig. 3E).

SlsA is an ortholog of the *P. alvei* S-layer protein SpaA, and carries an N-terminal SLH anchoring module followed by eight immunoglobulin-like domains, the architecture of the characterized S-layer proteins SbsC and Sap (Fig. 3D). Seven of its eight domains fall within three residues of the corresponding domain of SpaA. FLT43_RS14755 is likewise a tandem immunoglobulin-like array, but its SLH module lies at the opposite end (C-terminus) from every characterized structural S-layer protein. It is the homolog of *P. alvei* SlhA, an accessory SLH-domain surface protein of that organism involved in swarming and biofilm formation (20).

*slsA* is immediately downstream of FLT43_RS14755, and both are enclosed within a locus that also encodes *csaB* and the SCWP glycosyltransferase genes *tagA* and *tagO* homologs, with the same synteny found in *P. alvei* (Fig. 3E). Across the 26 surveyed genomes the locus is conserved, with *csaB*, *tagA*, *tagO* and the flanking housekeeping gene *fabZ* each above 97% nucleotide identity, while *slsA* falls to 67.7 to 84.6%. At the protein level, SlsA occurred as four distinct variants sharing 58 to 79% pairwise amino acid identity.

We identified the acid-extracted material as an S-layer based on its abundance, its SLH-mediated attachment to the cell wall, its homology to characterized *Bacillaceae* S-layer proteins, its release by standard low-pH glycine extraction, and the completeness of the blockage it imposes on a wall-binding probe.

In a whole-cell binding assay, after 5 min of acid treatment, the probe mGL-BDPt7 bound to the cell surface, as shown by its association with the cell pellet after washing (Fig. 3B). Binding was shown by the appearance of a band at the position of mGL-BDPt7 that corresponds to the computed mass of the expressed fusion, matches the probe-only control, and is absent from every preparation to which no probe was added (Fig. 3B). By contrast, the untreated pellet carried no distinguishable corresponding band. The same high-mass species seen in panel A are abundant in the untreated preparation and depleted after acid treatment.

### mGL-BDPt9 labels clinical isolates of 2 Paenibacillus species causing infant disease

We then investigated whether our labeling protocol could be effectively extended to other *P.* thiaminolyticus strains. Upon glycine-HCl treatment, purified mGL-BDPt9 successfully labeled all 6 tested strains spanning both clinical and non-clinical origins, including Mbale, Mbale2 and Mbale3, the 3 clinical isolates recovered from the cerebrospinal fluid of Ugandan infants with post-infectious hydrocephalus (33) (Fig. 4A–C), the reference strain ATCC 13023 (Fig. 4D–F), the species type strain NRRL B-4156 and NRRL NRS-1591 (Fig. S4).

**Figure 4.**
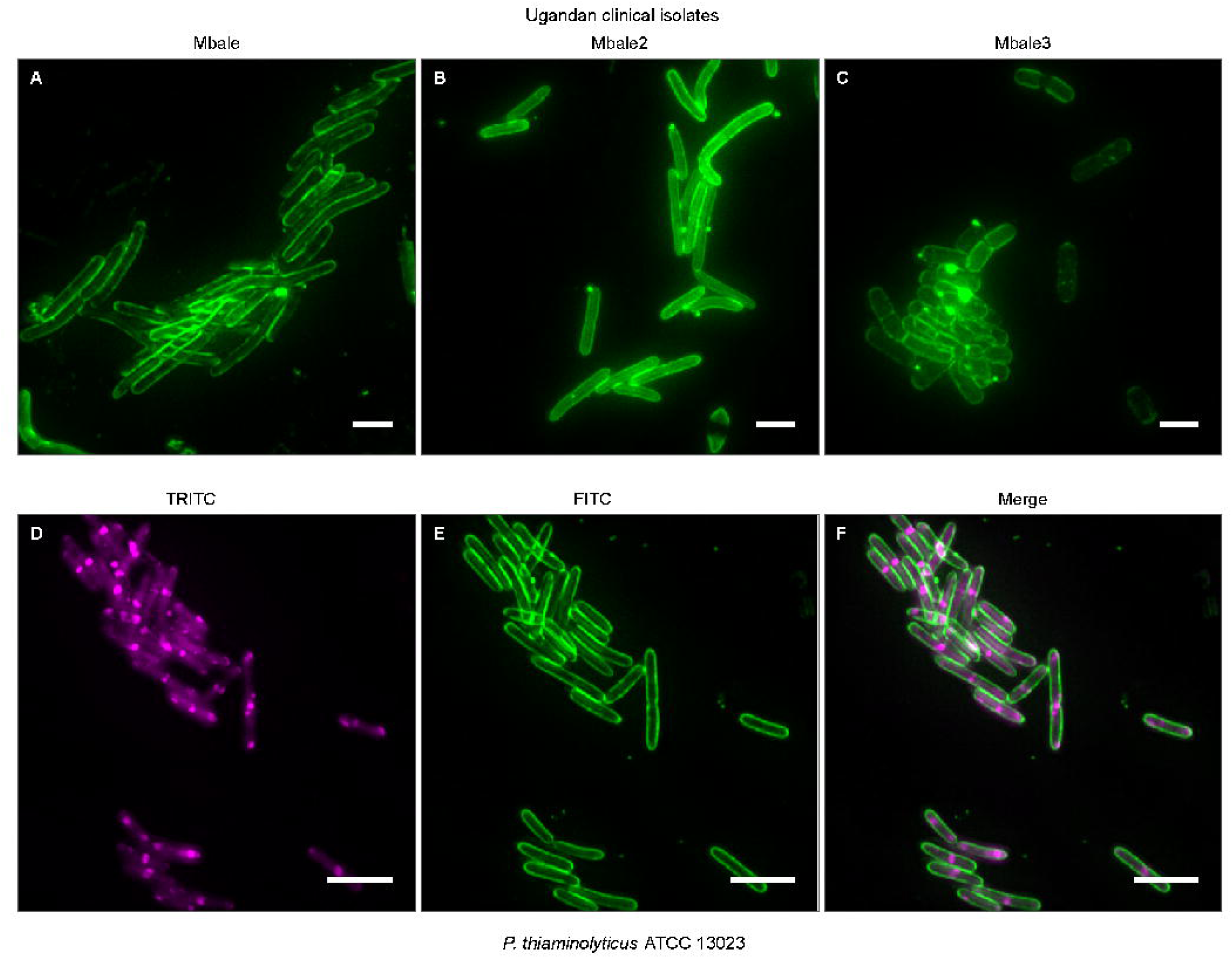
mGL-BDPt9 labels Ugandan clinical isolates of *Paenibacillus thiaminolyticus* and a reference strain. (Top) Three clinical isolates recovered from the cerebrospinal fluid of Ugandan infants with postinfectious hydrocephalus, namely Mbale (A), Mbale2 (B) and Mbale3 (C), treated with the glycine-HCl protocol, probed with purified mGL-BDPt9 and shown in the FITC channel. (Bottom) The reference strain *P. thiaminolytic*us ATCC 13023 was treated, counterstained with BacLight Red, and probed in the same way. Columns: TRITC channel (D), FITC channel (E), and the merge (F). Cells were fixed in 0.5% paraformaldehyde before treatment to stop cell motility. The two blocks were acquired at different magnifications: 60x for the clinical isolates and 100x for the reference strain. Scale bars: 5 µm.

In vegetative cells, the observed signal co-localized with the BacLight Red total-cell counterstain in the reference strain, outlining the counterstained cells and consistent with a target in the cell wall layer rather than the cytoplasm (Fig. 4D-F). The probes also variably outlined cells at various stages of sporulation, highlighting varied morphotypes that resembled stages described for other *Bacillus* and *Clostridium* species (Fig. S4). Interestingly, the same acid protocol enabled mGL-BDPt9 to label a *P. dendritiformis* strain recovered from an infected infant in the US, suggesting that it may carry a homologous surface barrier, similarly removed by acid wash (Fig. S5). Our lead probe therefore labels clinical isolates of *Paenibacillus* that are associated with infant disease, and the labeling similarly required prior envelope disruption.

After finding that the probe labeled clinical strains from two different *Paenibacillus* species, we asked how far its reach extends within the genus. Applied by quick-stain to 5 further species under the same conditions, mGL-BDPt9 stained *P. macerans* ATCC 8509, *P. polymyxa* ATCC 43865 and *P. barengoltzii* NR-36439, but could not stain *P. validus* (syn. *P. gordonae*) ATCC 29948 and *P. larvae* ATCC 9545 (Table 2). Because the envelope disruption protocol was not optimized for these species, we cannot rule out the possibility of labeling under modified protocols.

**Table 2.**
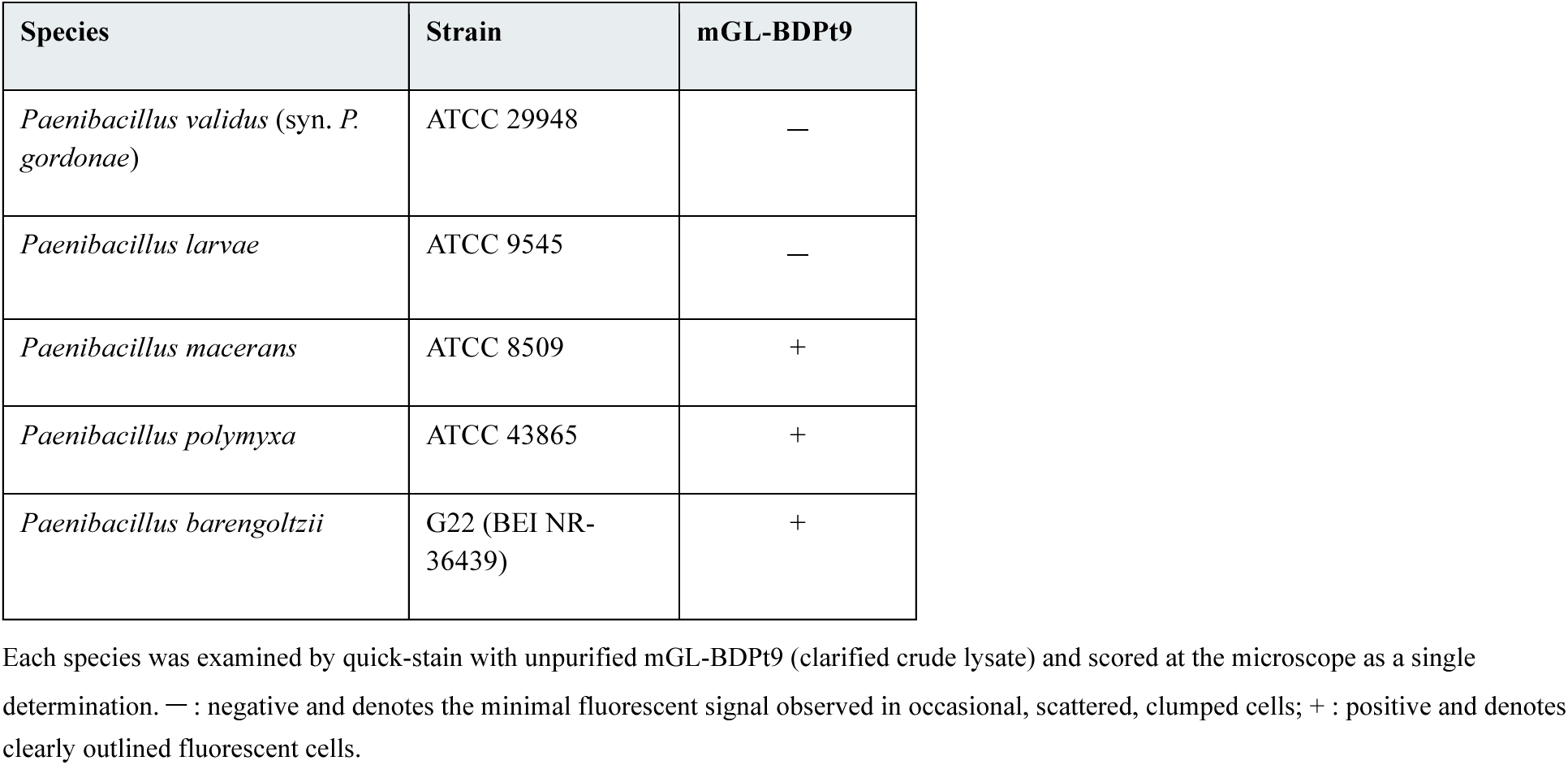
Labeling of five additional *Paenibacillus* species by mGL-BDPt9.

### mGL-BDPt9 distinguishes P. thiaminolyticus from 5 neonatal gram-positive sepsis pathogens in pairwise mixtures

Since the probe labeled other species within the genus, we tested its specificity against a panel of 5 gram-positive organisms that cause neonatal sepsis. After acid treatment, we mixed *P. thiaminolyticus* ATCC 13023 with *Streptococcus agalactiae* A909, *Staphylococcus aureus* NCTC-8325, *Streptococcus pyogenes* SF370, *Enterococcus faecalis* ATCC 29200, or *Listeria monocytogenes* HER 1184, then double-stained and imaged each mixture as a single preparation (Fig. 5). In every pairing, both species appeared in the counterstain channel, whereas only *P. thiaminolyticus* had a bright outline in the probe channel, showing that the probe could resolve our target species from the select panel of pathogenic organisms without prior isolation.

**Figure 5.**
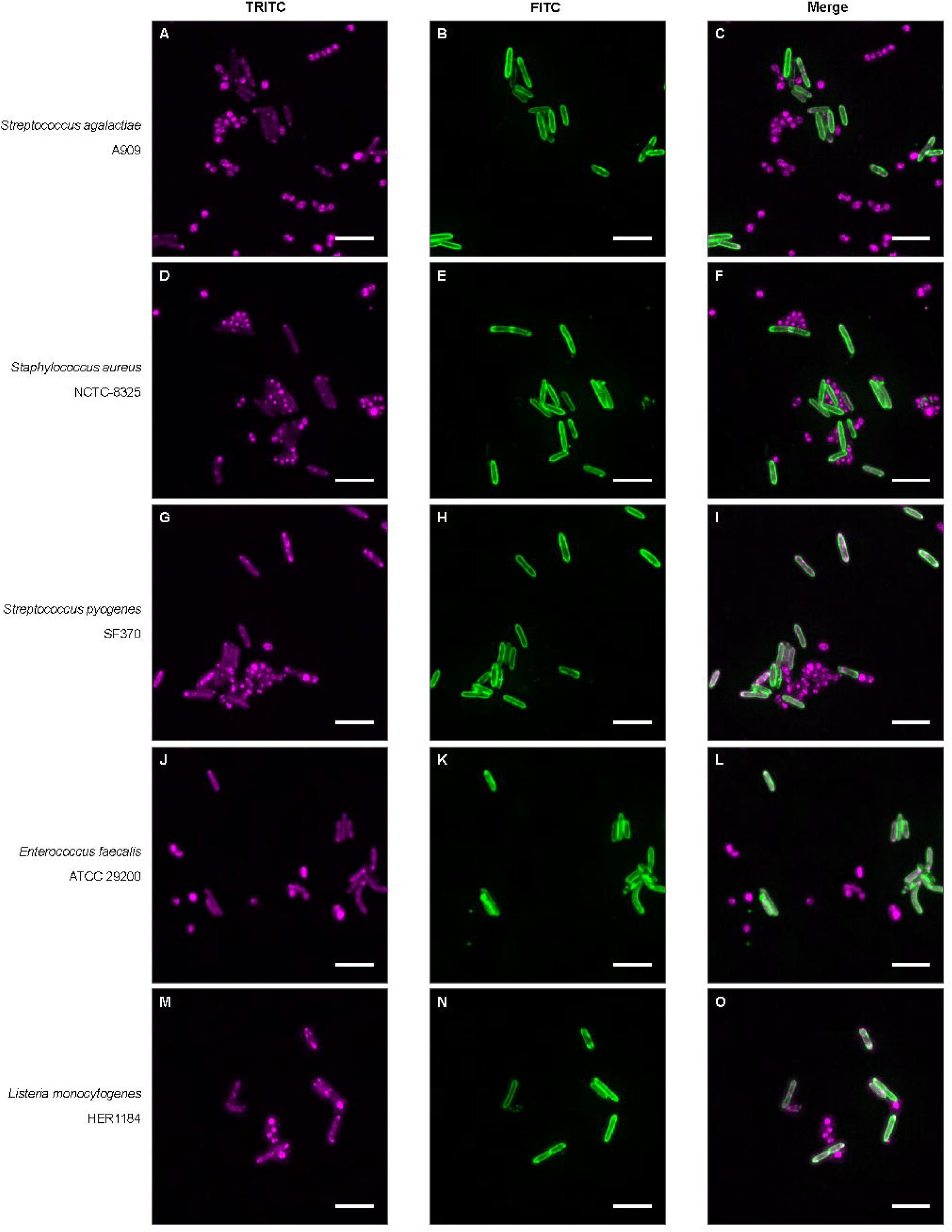
mGL-BDPt9 labels *P. thiaminolyticus* selectively in pairwise co-mixtures with gram-positive pathogens. *P. thiaminolyticus* ATCC 13023, treated with 0.2 M glycine-HCl pH 2 for 1 min, was mixed with each of 5 gram-positive species at equal optical density: *S. agalactiae* A909, *S. aureus* NCTC-8325, *S. pyogenes* SF370, *E. faecalis* ATCC 29200, and *L. monocytogenes* HER1184. Preparations were labeled with purified mGL-BDPt9, mounted without washing, and imaged at 100x. Phase contrast (left) and FITC (right) are shown for each pair. Rod-shaped cells with fluorescent outlines correspond to *P. thiaminolyticus*; cocci and shorter rods do not label. Scale bars: 10 um.

### mGL-BDPt9 resolves P. thiaminolyticus within a five-species suspension under the quick-stain protocol after flame fixation

The quick-stain protocol is the less resource-intensive of the two routes, and it had so far been applied to a single species. We tested it on a five-species suspension to assess whether the probe would still specifically resolve *P. thiaminolyticus*.

We prepared a suspension containing *E. coli DH5α*, *S. agalactiae*, *S. aureus*, *E. faecalis*, *and P. thiaminolyticus* at matched optical densities, flame-fixed it, and stained it with purified mGL-BDPt9 without washing (Fig. 6A to 6D).

**Figure 6.**
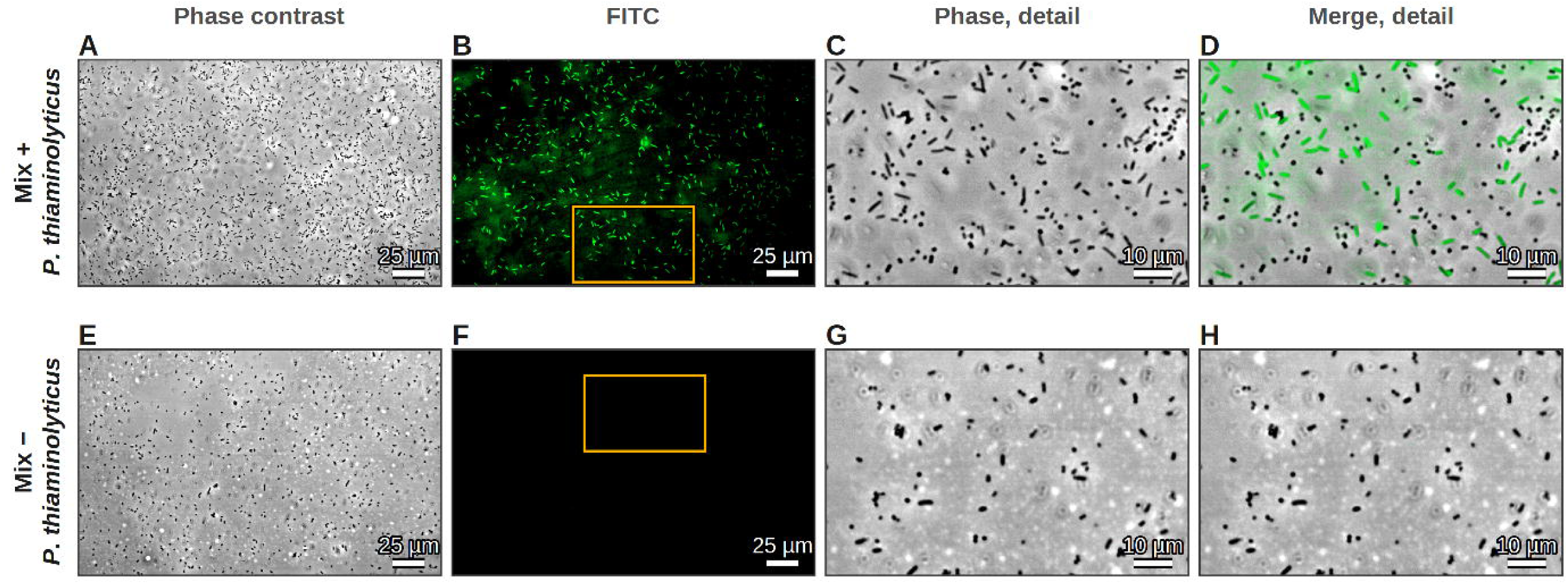
mGL-BDPt9 detects *Paenibacillus thiaminolyticus* in a five-species mixture after flame fixation. (A–D) A suspension *of P. thiaminolyticus* ATCC 13023, *Streptococcus agalactiae* A909, *Staphylococcus aureus* NCTC-8325, *Enterococcus faecalis* ATCC 29200 and *Escherichia coli* DH5α, flame-fixed and stained with purified mGL-BDPt9. (E–H) The same suspension prepared without *P. thiaminolyticus*, stained in the same session. Columns: phase contrast and FITC of the whole field, then phase contrast and the merged image of the boxed detail region. Preparations were not washed. Scale bars: 25 μm (whole field) and 10 μm (detail).

Labeling resolved a distinct homogeneous subpopulation of the mixture, representing approximately one cell in five in the field, uniform in shape and size and resembling *P. thiaminolyticus* in Fig. 2. The same suspension prepared without the target, stained in the same session, carried no labeled cells (Fig. 6E to 6H), including the gram-negative rod *E. coli* DH5α, indicating that labeling was therefore contingent on the target’s presence. Neither the mixture nor the target alone gave a signal when the probe was omitted (Fig. S6). The probe also detected its target in a multispecies mixture when applied as unpurified lysate (Fig. S7), under a protocol requiring only a flame and a slide.

## DISCUSSION

In this study, we developed the first affinity reagents for direct labeling of *Paenibacillus thiaminolyticus*. Five cell wall-binding probes labeled the pathogen only after acid treatment removed a barrier on the cell surface, a previously undescribed S-layer.

We identified SlsA as the major structural component of this surface barrier based on biochemical and genomic features characteristic of bacterial S-layers. SlsA is the ortholog of the characterized *Paenibacillus alvei* S-layer protein SpaA, is highly abundant at the cell surface, and is released by an acidic wash, a standard method for extracting S-layer proteins (22, 37). Untreated cell suspensions remained largely unlabeled by our probes but became uniformly outlined after SlsA was stripped from the cells. This binary labeling pattern is consistent with SlsA forming a continuous barrier on the cell surface. This structure appears to be widespread in *P. thiaminolyticus*, as indicated by the conservation of the secondary cell wall polymer (SCWP)- and S-layer-associated locus encoding SlsA across every sequenced genome surveyed (Fig. 3E, Table S2), and by consistent reproduction of the same labeling phenotype in every strain tested (Fig. 4, Fig. S4).

The dependence of labeling on SlsA removal, together with the homology between its SLH anchoring module and the binding domains of our 5 active probes, suggests that SlsA and the probes recognize a common cell-wall ligand. Structural work on *P. alvei* SpaA places SLH binding on the terminal pyruvylated monosaccharide of an SCWP (19, 34), making this a plausible candidate. However, some SLH modules have been shown to target cell wall components other than SCWPs, such as *Clostridium thermocellum* SdbA, which binds peptidoglycan directly (38), so further studies are needed to characterize the specific ligand recognized by our probes. The most parsimonious model is that SlsA occupies the ligand while anchoring to the cell wall, and that acid extraction frees it, allowing probe binding. Additionally, molecular sieving by the S-layer barrier may also contribute to restricting probe access to the wall, although the present experiments cannot distinguish between the two mechanisms (39, 40).

Our probes carry a single SLH domain at the C-terminus (Fig. 1), unlike the most studied SLH modules, which occur as 3 tandem repeats at the N-terminus (41). At the time we selected these candidates, no SLH domains had been reported as possible endolysin CBDs (42, 43). Since then, LysBT1 has been described independently in a prophage of *Brevibacillus thermoruber* (44), providing independent confirmation of this unusual architecture. Our study identified 4 novel phage endolysins with the same architecture, plus a closely related cell-wall hydrolase. LysBT1’s single C-terminal SLH domain was proposed to assemble into a homotrimer, forming a binding site similar to the three-fold modules of S-layer proteins (19, 44), so the single domain in our probes may behave similarly. Alternatively, it could bind in its monomeric form, as *P. alvei* SlhA, which retains residual binding to cell wall sacculi when 2 of the 3 SLH domains in its module are deleted (20). Interestingly, mGL-BDPt6, the only candidate carrying the canonical 3 repeats, did not label. SLH repeat number alone therefore did not predict binding in this system.

In organisms whose cell wall is concealed by a surface structure, intact-cell binding assays can misdirect the selection of CBD-based affinity reagents. Pretreatment has enabled or improved CBD labeling in several other organisms, including *Clostridium botulinum*, *Clostridioides difficile*, *Bacillus cereus*, and *Mycobacterium smegmatis* (21, 22, 44, 45). Had we limited our screening to intact cells, we would have discarded all 5 binders reported here, including mGL-BDPt9. The pretreatment itself was a single minute at pH 2, which left the cells motile and their gross morphology, rather astoundingly, unchanged. A few seconds of flame on a slide similarly enabled probe labeling, providing an alternative route. These findings show that the S-layer can act as a substantial barrier on the intact cell, but its removal is simple and adds little complexity to detection, capture, and research applications that do not require the native surface to be preserved. The 5 functional probes were developed from publicly deposited genome sequences and screened as crude recombinant lysates, providing a rapid route to affinity-reagent development for an emerging organism lacking established detection reagents.

*Paenibacillus* can be identified by MALDI-TOF only after successful isolation, thus inheriting every limitation of the preceding culture step (8). A probe directly recognizing the organism could move detection upstream of culture, where failure to recover the bacterium has made identification very challenging. Among 37 neonates with paenibacilliosis, routine aerobic blood cultures grew an organism in only 5 cases, none of them *Paenibacillus*, and no cerebrospinal fluid (CSF) culture grew bacteria in the local laboratory (6). In this study, we performed the labeling protocol exclusively on cultured cells, so our data first support its use to confirm cultured isolates, for which the acid step is trivial to incorporate. Although we did not test performance in clinical specimens, applying an adapted labeling protocol to CSF and other biological samples could have greater clinical utility, bypassing culturing and enabling rapid detection in a normally sterile compartment. Lysin-derived CBDs can recognize their cell wall ligands with nanomolar affinity (46) and, when immobilized on paramagnetic beads, have been used to enrich target organisms from mixed cultures and complex matrices (13, 47). Coupling such a capture step to a readout that does not require microscopy, such as lateral flow, could extend the approach to settings where microscopy is impractical.

In our preliminary characterization studies, mGL-BDPt9 labeled 4 additional *Paenibacillus* species, including a clinical isolate of *P. dendritiformis* recovered from an infant in the United States (Table 2, Fig. S5) (8), another cause of human paenibacilliosis (7, 9, 10), suggesting future usefulness for targeting human-associated *Paenibacillus* infections more broadly. None of the organisms we tested outside the genus, including common neonatal pathogens, were positive, either in pairwise co-mixture with the target (Fig. 5) or in the 5-species suspension, in which no labeling occurred without *P. thiaminolyticus* (Fig. 6, Fig. S7). The only two binding-domain probes previously reported in this genus were based on domains unrelated to the SLH module, and neither labeled the other’s host species (15, 16). Besides *Paenibacillus,* we did not test any other spore-forming members of the Bacillales, whose species largely rely on SLH domains to anchor surface proteins to pyruvylated wall polymers (18, 19). The probe’s labeling capacity in *Bacillus* and related spore-forming bacteria should therefore be assessed. Notably, the conditions required to expose the ligand may themselves be species-dependent, so tailoring the S-layer-stripping treatment offers a route to increasing specificity or to broadening coverage within the genus.

Beyond probe development, the identification of an S-layer contributes a major surface structure to the known biology of *P. thiaminolyticus*. S-layers form the structural core of a wider surface architecture, with many other proteins anchored to the wall through the same SLH-mediated mechanism (18, 48). The genome of *Bacillus anthracis* encodes 23 other SLH-domain proteins, including the adhesin BslA, which is required for penetration of the blood-brain barrier but dispensable for bacteremia in mice (49, 50). Similarly, in addition to SlsA, we found 26 SLH-containing proteins of undetermined function encoded in the *P. thiaminolyticus* reference genome, and we directly detected 3 of them in the acid surface extract. Future studies are needed to investigate the role of the *P. thiaminolyticus* S-layer and its associated proteins in invasion and central nervous system infection.

## ACKNOWLEDGMENTS

This research was supported by the Stavros Niarchos Foundation (SNF) as part of its grant to the SNF Institute for Global Infectious Disease Research at The Rockefeller University. The funder had no role in study design, data collection and interpretation, or the decision to submit the work for publication. We thank Henrik Molina of the Proteomics Resource Center at The Rockefeller University (RRID:SCR_017797) for LC-MS/MS acquisition and database searching. We thank Alison North and Ivan Suarez from the Rockefeller University’s Bio-Imaging Resource Center, RRID:SCR_017791, for help with imaging and image analysis. The following reagent was obtained through BEI Resources, NIAID, NIH: Paenibacillus barengoltzii, Strain G22, NR-36439.

