## Supplemental information for "Removing an S-layer enables endolysin-derived probes to detect *Paenibacillus thiaminolyticus* in mixed bacterial populations"

#### SUPPLEMENTARY METHODS

*Plasmid construction*

Binding-domain fragments were amplified from the parent constructs and assembled into a pET14b-based mGreenLantern fusion vector by in vivo assembly, using primers carrying 25-nt homologous tails. Vector PCR product was digested with DpnI (1 h, 37 °C) before assembly. Assemblies were transformed into *Escherichia coli* DH5α, screened by colony PCR on 1% agarose gels, and confirmed by Sanger sequencing. Confirmed plasmids were transformed into *E. coli* BL21-AI.

*Protein expression and purification*

Transformed BL21-AI were grown in Terrific Broth at 37 °C with shaking at 200 rpm to $OD_{600}$ 0.5, cooled on ice, induced with 0.2% (wt/vol) arabinose and 0.5 mM IPTG, and grown 18 h at 18 °C. Cells were harvested, washed twice in PBS and stored at −80 °C. For binding experiments, pellets were resuspended in cold PBS, sonicated on ice (5 min, 50% amplitude, 10 s on / 10 s off), clarified by centrifugation (20,000 × g, 15 min, 4 °C) and filtered. This clarified sonicate is the crude lysate referred to throughout. For purified preparations, clarified lysate was applied to Ni-NTA agarose, washed, eluted in 300 mM imidazole, dialyzed into Buffer A, concentrated and quantified by BCA assay.

*Expression and solubility screening (Fig. S1)*

Total-lysate and clarified-supernatant samples of all eleven fusions were resolved by SDS-PAGE as described in Materials and Methods and stained with SimplyBlue SafeStain. All eleven were detected in the crude lysate. The soluble fraction of mGL-BDPt8 was visibly depleted relative to the others.

*Eleven-construct binding screen (Fig. S2, Table 1)*

Clarified crude preparations of all eleven fusions were applied to *P. thiaminolyticus* ATCC 13023, first to intact cells and then to cells treated with 0.2 M glycine-HCl pH 2.0 for 1 min. The operator examined each field and scored it positive or negative against background. Micrographs of the constructs that were labeled are shown in a shared display window.

*Glycine-HCl dose-response (Table S3)*

Two independently prepared biomass sources, one scraped from agar colonies and one harvested from an overnight broth culture, were treated with Buffer A or with 0.2 M glycine-HCl at pH 4, 3 or 2 for 1 or 5 min, labelled with mGL-BDPt9 and scored at the microscope. Labelling and cell morphology were recorded for each condition and biomass source.

*Additional strains and species (Table 2, Fig. S4, Fig. S5)*

*P. thiaminolyticus* NRRL B-4156 and NRS-1591, five additional *Paenibacillus* species and *P. dendritiformis* were prepared as described in the main methods and examined with mGL-BDPt9, each before and after envelope disruption. Species and strains are listed in Table S4.

*No-probe and crude-lysate controls (Fig. S6, Fig. S7)*

For no-probe controls, *P. thiaminolyticus* alone and the five-species suspension were carried through the full preparation with Buffer A in place of probe, and two fields of each were acquired at the settings used for the corresponding experimental fields. For the crude-lysate field, the five-species suspension was overlaid with clarified crude lysate of *E. coli* expressing mGL-BDPt9 in place of purified probe.

*MS peptide coverage (Fig. S3)*

Peptide-spectrum matches for each excised band were mapped onto the SlsA precursor sequence and plotted against the predicted domain boundaries. Coverage is reported for the mature sequence after signal-peptide cleavage.

*Locus alignment (Fig. 3E)*

Loci were aligned as described in Materials and Methods. The same settings were used for the 26-genome and the six-genome alignments. Sequence direction was determined automatically.

*Deconvolution*

DeltaVision stacks were bleach- and z-line-corrected before deconvolution, and deconvolved by three-dimensional iterative constrained deconvolution against a measured optical transfer function rescaled to each emission wavelength. Deconvolution trims a border, and exported panels are the valid region of the acquired field.

*Genome assemblies and accessions (Table S1, Table S2)*

Accessions are given in GenBank (GCA) format throughout, because one assembly, that of strain Mbale3, has no RefSeq equivalent. Six of the 26 accessions represent the type strain, deposited variously as NRRL B-4156, NRRL B-04156, NBRC 15656, DSM 7262 and MGYG-HGUT-02490.

### SUPPLEMENTARY FIGURES AND TABLES

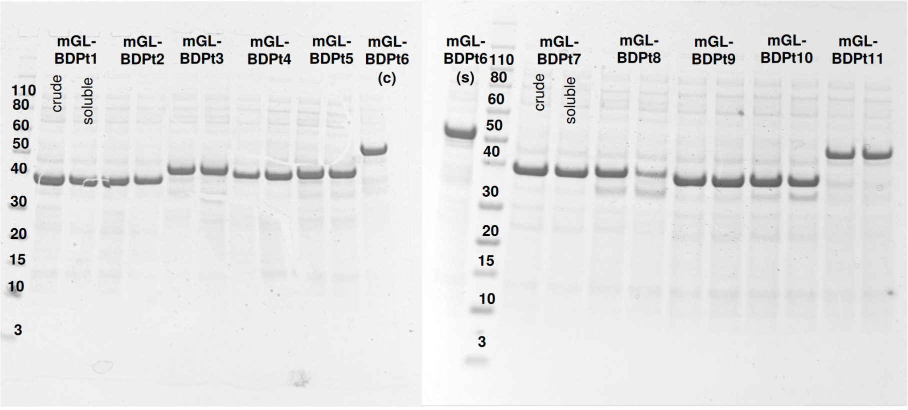


**Figure S1. Expression of the eleven mGreenLantern binding domain fusions in *Escherichia coli***
Crude lysate and the soluble fraction of each fusion, resolved by SDS-PAGE and stained with Coomassie blue. Each fusion appears as a band between 36 and 50 kDa. Two gels are shown side by side. The soluble fraction of mGL-BDPt8 is depleted relative to its crude lysate, and mGL-BDPt6 runs above the other ten.

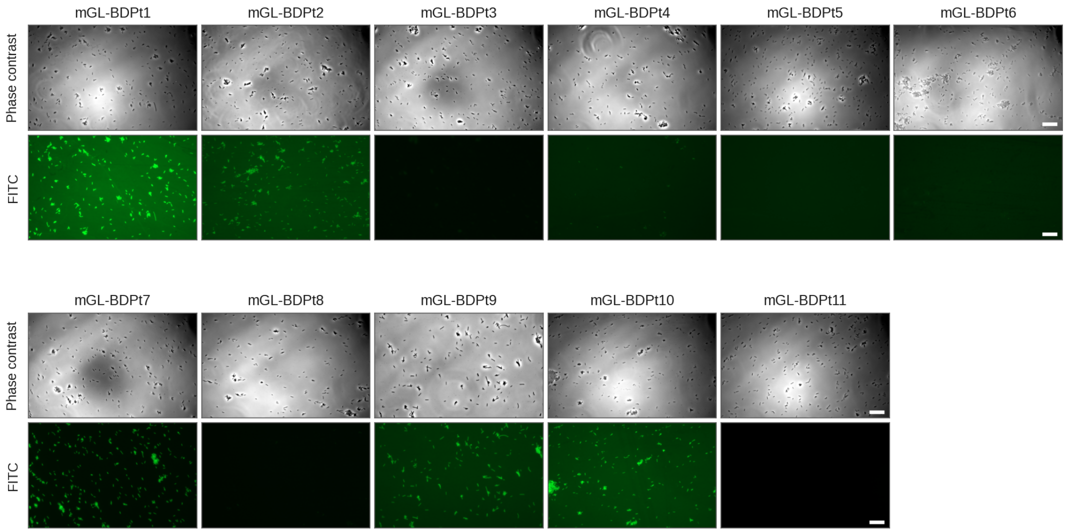


**Figure S2. Labelling of *Paenibacillus thiaminolyticus* ATCC 13023 by eleven mGreenLantern binding domain fusions**

Cells treated with 0.2 M glycine-HCl pH 2 for 1 min were incubated with clarified crude lysate of each fusion and imaged at 40x in phase contrast and in the FITC channel. One shared linear FITC display window is applied to every panel, so panels may be compared directly for the presence and absence of signal. The phase contrast channel is scaled per panel. mGL-BDPt1, mGL-BDPt2, mGL-BDPt7, mGL-BDPt9 and mGL-BDPt10 labelled the cells. Scale bars: 25 um.

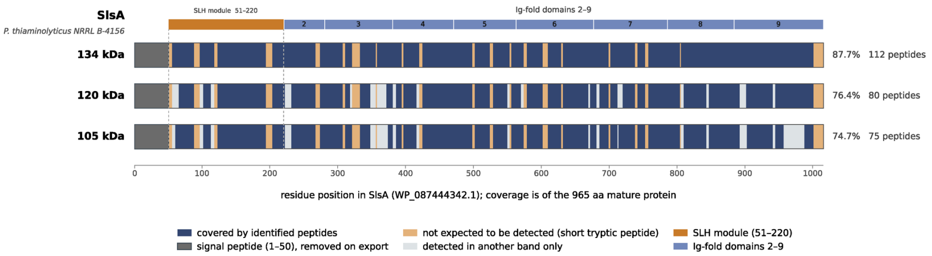


**Figure S3. LC-MS/MS peptide coverage of SlsA in the three acid-extractable high-molecular-mass bands**
Peptides assigned to SlsA (WP_087444342.1) following LC-MS/MS analysis of the apparent 134-, 120-, and 105-kDa bands excised from Fig. 3A are mapped onto the SlsA sequence. The predicted 1-50 signal peptide, which is removed during export, is hatched; the SLH anchoring module (residues 51-220) and eight predicted Ig-like domains are indicated above. Dark blue denotes sequence covered by peptides identified in the indicated band; amber indicates *in silico*-predicted tryptic peptides shorter than the 6-residue detection limit that were not detected. Some peptides from this class were detected within longer, partially digested peptides; gray indicates sequence covered in at least one of the other excised bands but not the indicated band. Sequence coverage of the 965-aa mature protein was 87.7% (112 peptides), 76.4% (80 peptides), and 74.7% (75 peptides) for the apparent 134-, 120-, and 105-kDa bands, respectively.

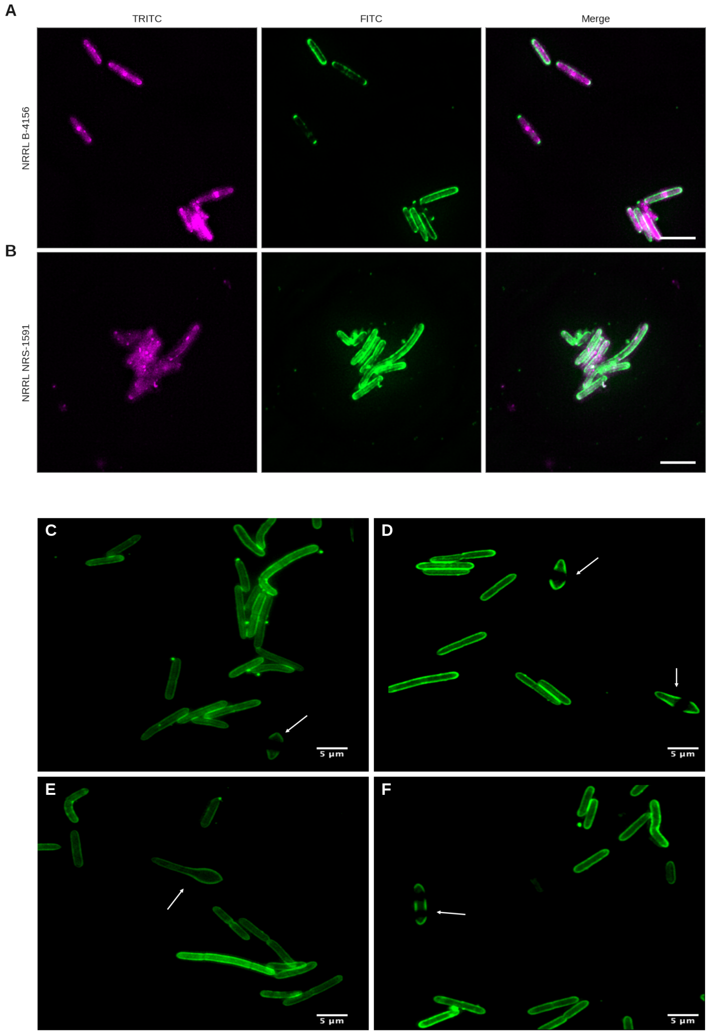


**Figure S4. mGL-BDPt9 labels two further *Paenibacillus thiaminolyticus* strains and labels sporulating cells**
(A, B) NRRL B-4156 and NRRL NRS-1591, counterstained with BacLight Red and probed with purified mGL-BDPt9. Columns: TRITC, FITC and the merged image. Scale bars: 5 um. (C-F) *P. thiaminolyticus* ATCC 13023 after prolonged culture, probed with mGL-BDPt9 and shown in the FITC channel. Arrows mark sporulating cells, in which the probe outlines the mother cell and the forespore. Scale bars: 5 um.

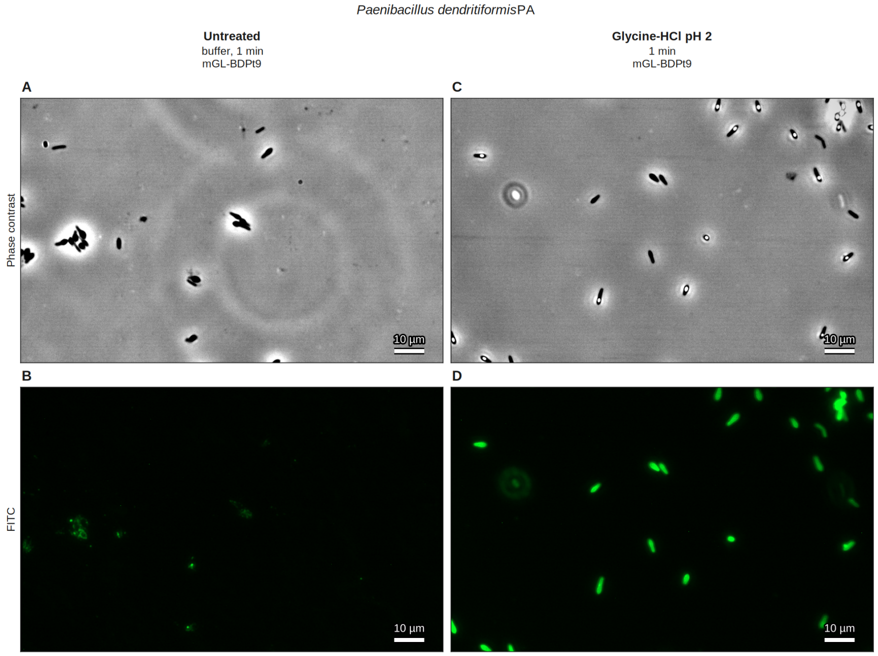


**Figure S5. Envelope disruption relieves a barrier to mGL-BDPt9 labeling of *Paenibacillus dendritiformis***

Phase contrast (A, C) and the FITC channel (B, D) of the same field. (A, B) Cells were kept in Buffer A for 1 min and incubated with purified mGL-BDPt9. (C, D) Cells treated with 0.2 M glycine-HCl pH 2 for 1 min and incubated with mGL-BDPt9. The heterologous control fusion mGL-BDCp17 was scored negative in both untreated and treated cells and is not shown. Scale bars: 10 um.

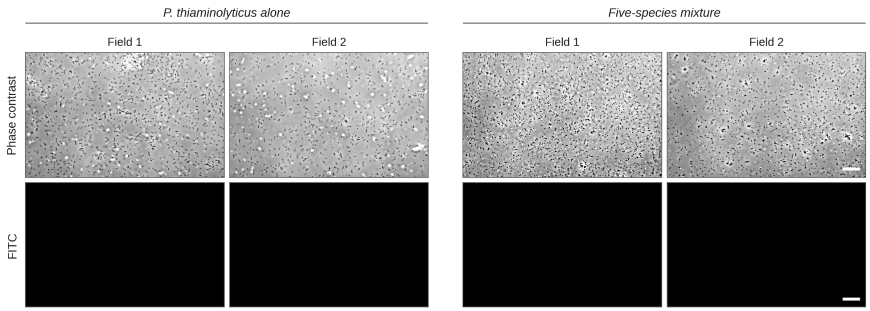


**Figure S6. No labelling without probe**
(A-D) *P. thiaminolyticus* ATCC 13023 alone, and the five-species suspension of Fig. 6, each flame-fixed and treated with buffer in place of mGL-BDPt9. Two fields each, imaged in the same session and at the same settings as Fig. 6 and displayed in the windows of that figure: phase contrast 44159-60378, FITC 9776-17588 counts. Cells are present in phase contrast in every field and no signal is seen in the FITC channel. Scale bar: 25 um.

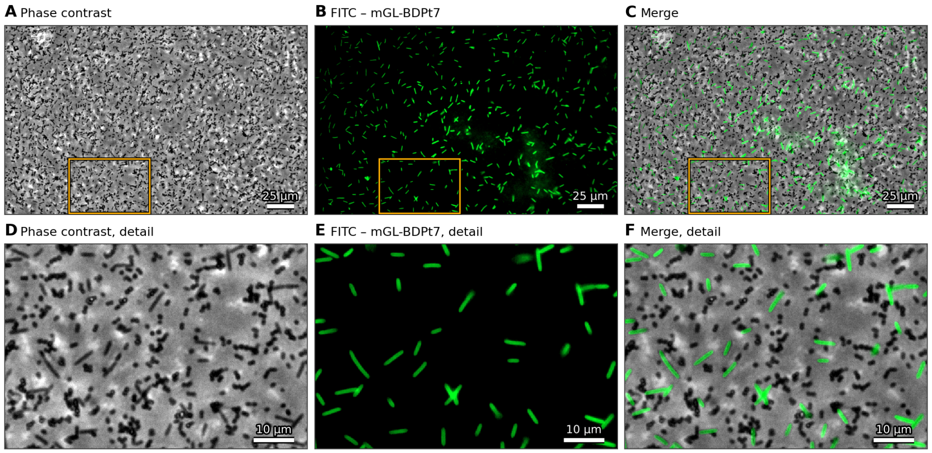


**Figure S7. mGL-BDPt9 in unpurified lysate resolves *Paenibacillus thiaminolyticus* within a five-species suspension**

A suspension of *P. thiaminolyticus* ATCC 13023 mixed with *Escherichia coli* BL21, *Streptococcus agalactiae* A909, *Enterococcus faecalis* ATCC 29200 and *Listeria monocytogenes* HER1184 at matched optical densities was flame-fixed on a slide, probed with unpurified mGL-BDPt9 overexpression lysate, and imaged without washing. This five-species panel differs from that of Fig. 6, in which *Staphylococcus aureus* NCTC-8325 takes the place of *L. monocytogenes*. (A-C) The whole field in phase contrast, the FITC channel and the merge. Gold rectangles mark the regions shown enlarged in (D-F). Rod-shaped cells carrying probe signal are interspersed with unlabelled cells of the other morphologies present. Scale bars: 25 um (A-C) and 10 um (D-F).

#### Table S1

| Candidate | Source strain | Isolation source | Assembly accession | Length (aa) | Mass (kDa) | pI | Domain architecture (residues) | Cloned binding domain | mGL-BDPt binding |
|---|---|---|---|---|---|---|---|---|---|
| LysPt1 | BO5 | Soil, Moscow | GCA_003591545.1 | 220 | 24.76 | 7.72 | Amidase_2 22–143 · SLH 163–220 | 167–220 | yes |
| LysPt2 | BO5 | Soil, Moscow | GCA_003591545.1 | 221 | 24.45 | 7.33 | Amidase_2 22–143 · SLH 163–221 | 163–221 | yes |
| LysPt3 | BO5 | Soil, Moscow | GCA_003591545.1 | 459 | 50.48 | 4.41 | Cu_amine_oxidN 3–108 · AMIN 164–246 · Amidase_3 284–453 | 6–113 | no |
| LysPt4 | BO5 | Soil, Moscow | GCA_003591545.1 | 241 | 26.55 | 10.17 | SH3_3 13–82 · NLPC_P60 95–240 | 17–82 | no |
| LysPt5 | NRRL B-4156 | Feces, Japan | GCA_007066085.1 | 331 | 36.69 | 4.99 | Peptidase_M23 208–298 | 49–139 | no |
| LysPt6 | Mbale | CSF, Uganda | GCA_007066225.2 | 507 | 56.58 | 4.51 | SLH 5–68 · SLH 70–128 · SLH 129–191 · Glyco_hydro_18 177–503 | 6–189 | no |
| LysPt7 | Mbale | CSF, Uganda | GCA_007066225.2 | 420 | 45.39 | 8.21 | Amidase_3 4–177 · Glucosaminidase 235–360 · SLH 363–420 | 363–420 | yes |
| LysPt8 | NRRL B-24730 | Environ., USDA | GCA_026794135.1 | 207 | 23.18 | 8.71 | LD_TPASE 19–128 · PG_binding_1 148–201 | 145–207 | no |
| LysPt9 | PATH554 | Feces, China | GCA_030169715.1 | 396 | 43.58 | 9.32 | Amidase_2 23–144 · SLH 340–396 | 340–396 | yes |
| LysPt10 | PATH554 | Feces, China | GCA_030169715.1 | 220 | 24.75 | 7.70 | Amidase_2 21–143 · SLH 163–220 | 161–220 | yes |
| LysPt11 | Mbale3 | CSF, Uganda | GCA_029532855.1 | 524 | 58.08 | 4.38 | Cu_amine_oxidN 28–105 · SH3_3 133–183 · Glyco_hydro_18 208–521 | 116–191 | no |

**Table S1. The eleven candidate cell-wall hydrolases of *Paenibacillus thiaminolyticus***
Candidates were mined from publicly available genome assemblies classified as *P. thiaminolyticus*. Length, molecular mass and isoelectric point were computed from the mature sequence using Geneious. Domains were annotated with InterProScan. The cloned binding-domain boundaries are those used to construct the mGreenLantern fusions and are drawn as rules in Fig. 1. Labeling results are those of Table 1.

#### Table S2

| Assembly name | Strain | Assembly accession |
|---|---|---|
| ASM359154v1 | BO5 | GCA_003591545.1 |
| DSM_7262 | DSM 7262 | GCA_945318275.1 |
| IPM_PthMbale_MinION | Mbale | GCA_007066225.2 |
| ASM2855441v1 | Mbale2 | GCA_028554415.1 |
| ASM2953285v1 | Mbale3 | GCA_029532855.1 |
| UHGG_MGYG-HGUT-02490 | MGYG-HGUT-02490 | GCA_902387705.1 |
| ASM400100v1 | NBRC 15656 | GCA_004001005.1 |
| 57566_C02 | NCTC11027 | GCA_900454555.1 |
| ASM2679767v1 | NRRL B-04156 | GCA_026797675.1 |
| ASM2679731v1 | NRRL B-14604 | GCA_026797315.1 |
| ASM2679725v1 | NRRL B-14605 | GCA_026797255.1 |
| ASM2679729v1 | NRRL B-14607 | GCA_026797295.1 |
| ASM2679726v1 | NRRL B-14608 | GCA_026797265.1 |
| ASM2679723v1 | NRRL B-14609 | GCA_026797235.1 |
| ASM2679721v1 | NRRL B-14610 | GCA_026797215.1 |
| ASM2679719v1 | NRRL B-14611 | GCA_026797195.1 |
| ASM2679717v1 | NRRL B-14612 | GCA_026797175.1 |
| ASM2679715v1 | NRRL B-14613 | GCA_026797155.1 |
| ASM2679713v1 | NRRL B-14625 | GCA_026797135.1 |
| ASM2679413v1 | NRRL B-24730 | GCA_026794135.1 |
| ASM706608v1 | NRRL B-4156 | GCA_007066085.1 |
| ASM216185v1 | NRRL B-4156 | GCA_002161855.1 |
| ASM3580367v1 | NRS-1444 | GCA_035803675.1 |
| ASM3580363v1 | NRS-1591 | GCA_035803635.1 |
| ASM3016971v1 | PATH554 | GCA_030169715.1 |
| ASM2822677v1 | SY20 | GCA_028226775.1 |

**Table S2. Genome assemblies of *Paenibacillus thiaminolyticus* surveyed for cell-wall hydrolase candidates**
All 26 assemblies classified as *P. thiaminolyticus* and publicly available at the time of screening, representing 21 distinct strains. Six accessions are of the type strain, deposited variously as NRRL B-4156, NRRL B-04156, NBRC 15656, DSM 7262 and MGYG-HGUT-02490. Accessions are given in GenBank format.

#### Table S3

| Condition | pH | Duration | Labelling (both biomass sources) | Morphology, Agar-scraped (colonies) | Morphology, Broth-harvested (overnight) | Note |
|---|---|---|---|---|---|---|
| 50 mM HEPES, 140 mM NaCl, pH 7.4 | 7.4 | -- | Very poor; labelling of scattered cells, often in clumps and chains | Long, often filamentous chains, frequently in clumps; small motile bacilli | Medium-length chains, some in clumps; small motile bacilli | untreated control |
| 0.2 M glycine-HCl, pH 4 | 4 | 1 min | Poor; slight improvement over the neutral-pH control, hard to call | Long, often filamentous chains, frequently in clumps; small motile bacilli | Medium-length chains, some in clumps; small motile bacilli | |
| 0.2 M glycine-HCl, pH 3 | 3 | 1 min | Some staining, some cells | Long, often filamentous chains, frequently in clumps; small motile bacilli | Medium-length chains, some in clumps; small motile bacilli | |
| 0.2 M glycine-HCl, pH 2 | 2 | 1 min | Best condition tested; all cells in frame | Long, often filamentous chains, frequently in clumps; small motile bacilli | Medium-length chains, some in clumps; small motile bacilli | adopted as standard |
| 0.2 M glycine-HCl, pH 2 | 2 | 5 min | No difference discernible by eye from 1 min | Long, often filamentous chains, frequently in clumps; small motile bacilli | Medium-length chains, some in clumps; small motile bacilli | |
| 0.2 M glycine-HCl, pH 3 | 3 | 5 min | No difference discernible by eye from 1 min | Long, often filamentous chains, frequently in clumps; small motile bacilli | Medium-length chains, some in clumps; small motile bacilli | |

**Table S3. Labelling of Paenibacillus thiaminolyticus by mGL-BDPt9 across a glycine-HCl dose-response grid**
Two independently prepared biomass sources, one scraped from agar colonies and one harvested from an overnight broth culture, were treated at each condition and scored at the microscope. The two sources agreed at every condition, so one scoring column is given. Morphology was identical across all conditions within a source and is given once per source. The condition adopted for all subsequent work is marked.

#### Table S4

| Organism | Strain | Source | Medium | Culture conditions |
|---|---|---|---|---|
| *Paenibacillus thiaminolyticus* | ATCC 13023 | ATCC | TH | aerobic, 37 °C |
| *Paenibacillus thiaminolyticus* | NRRL B-4156 | USDA | TH | aerobic, 37 °C |
| *Paenibacillus thiaminolyticus* | NRS-1591 | USDA | TH | aerobic, 37 °C |
| *Paenibacillus thiaminolyticus* | Mbale | Schiff Lab, Yale | TH | aerobic, 37 °C |
| *Paenibacillus thiaminolyticus* | Mbale2 | Schiff Lab, Yale | TH | aerobic, 37 °C |
| *Paenibacillus thiaminolyticus* | Mbale3 | Schiff Lab, Yale | TH | aerobic, 37 °C |
| *Paenibacillus dendritiformis* | PA | Ericson's Lab, Penn State | TH | aerobic, 37 °C |
| *Paenibacillus validus* (syn. *P. gordonae*) | ATCC 29948 | Microbiologics | TH | aerobic, 37 °C |
| *Paenibacillus larvae* | ATCC 9545 | Microbiologics | BHI | aerobic, 37 °C |
| *Paenibacillus macerans* | ATCC 8509 | Microbiologics | TH* | aerobic, 37 °C |
| *Paenibacillus polymyxa* | ATCC 43865 | Microbiologics | TH | aerobic, 37 °C |
| *Paenibacillus barengoltzii* | G22 (BEI NR-36439) | BEI | TH | aerobic, 37 °C |
| *Staphylococcus aureus* | NCTC-8325 | RU collection | BHI | aerobic, 37 °C |
| *Streptococcus pyogenes* (M1) | SF370 | ATCC | BHI | aerobic, 37 °C |
| *Streptococcus agalactiae* | A909 | ATCC | BHI | aerobic, 37 °C |
| *Enterococcus faecalis* | ATCC 29200 | ATCC | BHI | aerobic, 37 °C |
| *Listeria monocytogenes* | HER1184 | RU collection | BHI | aerobic, 37 °C |
| *Escherichia coli* | BL21-AI | Thermo Scientific | LB | aerobic, 37 °C |
| *Escherichia coli* | DH5α | Thermo Scientific | LB | aerobic, 37 °C |

**Table S4. Bacterial strains used in this study.**
TH, Todd-Hewitt medium, 15 g/L agar; *TH with 0.1% Tween 80 to abate clumping. BHI, brain-heart infusion, 15 g/L agar. LB, Luria-Bertani, 15 g/L agar. *Escherichia coli* BL21-AI is the expression host for the mGreenLantern fusions and DH5-alpha the cloning strain, and were also included in the mixed population experiments.

#### Table S5

| Locus tag | Protein ID | Name | Annotation | Architecture | Mature kDa | Peptides | Cov. (%) | PSMs 105 kDa | Rank 105 kDa | PSMs 120 kDa | Rank 120 kDa | PSMs 134 kDa | Rank 134 kDa |
|---|---|---|---|---|---|---|---|---|---|---|---|---|---|
| FLT43_RS14730 | WP_087444338.1 | — | Ig-like domain-cont. protein | C-terminal SLH | 139.1 | 36 | 35.5 | 0 | nd | 0 | nd | 39 | 3/27 |
| FLT43_RS14755 | WP_087444343.1 | — | S-layer homology domain-cont. protein | C-terminal SLH | 130.4 | 48 | 45.8 | 13 | 6/22 | 8 | 4/10 | 81 | 2/27 |
| FLT43_RS14750 | WP_087444342.1 | SlsA | S-layer homology domain-cont. protein | N-terminal SLH | 106.6 | 112 | 87.7 | 1252 | 1/22 | 1369 | 1/10 | 3398 | 1/27 |
| FLT43_RS05020 | WP_087441867.1 | — | S-layer homology domain-cont. protein | N-terminal SLH | 98.3 | 45 | 60.9 | 68 | 2/22 | 0 | nd | 0 | nd |
| FLT43_RS14745 | WP_087444341.1 | orf7 | hypothetical protein | no SLH domain | 91.8 | 36 | 50.6 | 50 | 3/22 | 0 | nd | 0 | nd |

**Table S5. SLH-domain proteins identified by LC-MS/MS in the acid-extractable high-molecular-mass bands**.
Proteins carrying SLH domains, or co-eluting without an anchoring module (Orf7), detected alongside SlsA (FLT43_RS14750) in the acid-wash experiment of Fig. 3. Each was identified as a minor component of the indicated band. SlsA accounted for 81-97% of assigned spectra per band. PSMs, peptide-spectrum matches in the indicated band; nd, not detected. FLT43_RS14745 (orf7) carries no SLH domain at any detection threshold and is included because it co-released with the SLH-anchored proteins. Coverage is of the mature protein after removal of the predicted signal peptide.
